# Reliable single-cell perturbations explain and improve model performance

**DOI:** 10.64898/2026.08.11.744177

**Authors:** Xi Wang, Jack Kuipers, Florian Hugi, Randall J. Platt, Niko Beerenwinkel

## Abstract

Predicting single-cell transcriptional responses to perturbations is central to building the virtual cell, yet recent benchmarks show that simple baseline methods often outperform complex models, and model comparisons depend on the evaluation metric. Most studies assume that preprocessed RNA sequencing data are reliable ground truth for both training and evaluation. Here, we test this assumption by measuring the reliability of perturbations and their alignment with shared perturbation responses, classifying each perturbation as specific, shared, or unreliable. Among 7,170 perturbations from 29 datasets, 65% are unreliable, 11% shared, and 24% specific. Applying these quality labels to published benchmarks shows that model comparisons depend on perturbation quality. Training with reliable perturbations alone matches or outperforms full-data performance while using 55% of all training perturbations. Our framework also enables prospective experimental design: for most perturbations, a 28-cell pilot experiment accurately predicts how many cells a full screen needs to be reliable.

## 1 Introduction

Single-cell perturbation prediction is a crucial problem in functional genomics, drug discovery, and systems biology, and it sits at the heart of the recent virtual cell endeavors ^1,2^. Technologies such as Perturb-seq make it possible to measure transcriptional responses for hundreds to thousands of perturbations in a single experiment ^3–6^. As screening perturbations over all possible combinations of genes, cell states, doses, and time points is infeasible, the field utilizes deep-learning models to generalize from available perturbation measurements to unseen settings, in order to perform in silico perturbation experiments ^1,2,7^.

A growing literature has explored increasingly complex ways to represent cells and perturbations and to learn transformations from control to perturbed states ^8–12^. In parallel, transformer-based single-cell self-supervised foundation models have been proposed to learn network biology from millions of cells and have shown success in downstream fine-tuned tasks including perturbation modeling ^13–19^. Dedicated benchmarking platforms and competitions have also been created to accelerate progress toward predictive virtual cells ^1,2^.

Despite this rapid growth, two recent findings have cast doubt on the capabilities of current approaches. The first finding concerns model performance. Multiple independent studies, including large-scale benchmarks spanning dozens of methods and datasets, find that simple baselines, such as training-set means and linear predictors, match or outperform deep-learning and foundation models ^20–26^. The second finding concerns evaluation methodology. Studies report that different perturbations can induce a shared transcriptional shift relative to control, attributed mainly to biological programs such as stress response or cell-cycle arrest, so correlation-based metrics may reward prediction of the average perturbation effect rather than perturbation-specific effects ^27^. In addition, it has been shown that the choice of the evaluation metric alone can affect and even reverse performance-based model rankings ^28–32^. Together, these studies show that conclusions about perturbation-prediction models depend heavily on which aspect of the response is evaluated and how it is measured.

Beneath these findings lies an untested assumption, namely, that all single-cell perturbation data are suitable groundtruth targets for training and evaluating predictive models. The premise is increasingly consequential, as existing screens designed for biological discovery are being repurposed for model training and benchmarking. For each perturbation, the observed perturbed cells are a finite sample from the underlying population, and likewise for the control cells. The effect of the perturbation is thus estimated as the difference between the mean preprocessed expression profiles of the sampled perturbed and control cells. If this estimate is unstable across independent samples of cells under the same conditions due to noise, evaluation is limited by sampling and measurement noise, and training is based on an unstable target.

Here, we introduce a reliability framework, grounded in classical test theory ^33–35^, that makes data quality an explicit part of perturbation modeling. For each perturbation, we compute split-half reliability, the correlation between effect estimates from two independent halves of the cells, and combine it with a systematic-variation score to classify perturbations into the three data quality categories *specific* (reliable and perturbation-specific), *shared* (dominated by the shared response), and *unreliable* (dominated by noise).

We apply our framework to 29 preprocessed single-cell perturbation datasets from scPerturBench ^26^, covering 7,170 perturbations, and report three main findings. First, most perturbations are dominated by noise or by the shared response, with only 24% classified as specific. Second, these quality labels affect both evaluation and training. In published benchmarks, our framework shows that model comparisons depend on perturbation quality: nearly all methods score highest on shared perturbations, and restricting the evaluation to specific perturbations changes the top-ranked method in 5 out of 8 benchmark settings. In training, restricting training data to reliable perturbations recovers fulldata performance. Third, we derive a simple model linking reliability to cell count through the perturbation-specific per-cell signal-to-noise ratio. Because the number of cells required to reach a given target reliability scales inversely with this ratio, a small pilot experiment can separate perturbations that need fewer cells from those that require many more and those that are impractical due to the large number of cells required. We find that a pilot screen of 28 cells per perturbation accurately predicts the cell count required to be reliable for most perturbations, providing a basis for allocating sequencing effort across a full screen in experimental design.

## 2 Results

### 2.1 Reliable and specific perturbations are in the minority across 29 datasets

In single-cell perturbation studies, the response of a perturbation is usually quantified by the change in gene expression from the control to the perturbed cells using RNA sequencing. Most studies apply standardized preprocessing steps to the sequencing data which are then used as the ground truth for both model evaluation and training. Since gene expression measurements are noisy and different perturbations can induce similar transcriptional responses, we asked (*i*) whether the perturbation effect is reliable under resampling of cells, and (*ii*) how much of the measured response is shared across all perturbations in the dataset.

For each perturbation, we estimated measurement reliability *ρ* as the median Pearson correlation between two halves of its associated cells across 100 random splits, corrected for the reduced sample size using the Spearman–Brown formula. We quantified specificity *φ* as the squared cosine similarity between the perturbation’s effect vector and the mean response over cells in all reliable perturbations in the dataset ^27,33^.

We hypothesized that perturbations differ in whether their measured effects are reliable across sampled cells and specific to the perturbation rather than aligned with a response shared across the screen. To test this hypothesis, we classified each perturbation using the threshold 0.5 for both *ρ* and *φ* (Figure 1a–c). Each threshold marks a 1:1 variance ratio (signal to noise for *ρ*; perturbation-specific to shared for *φ*; Methods 3). We labeled perturbations with *ρ <* 0.5 as unreliable. Among the reliable (*ρ* ≥ 0.5), those with *φ* ≥ 0.5 were labeled shared because their response aligns with the transcriptional axis common to all perturbations in the dataset. The remainder were labeled specific, dominated by perturbation-specific signal.

**Figure 1:**
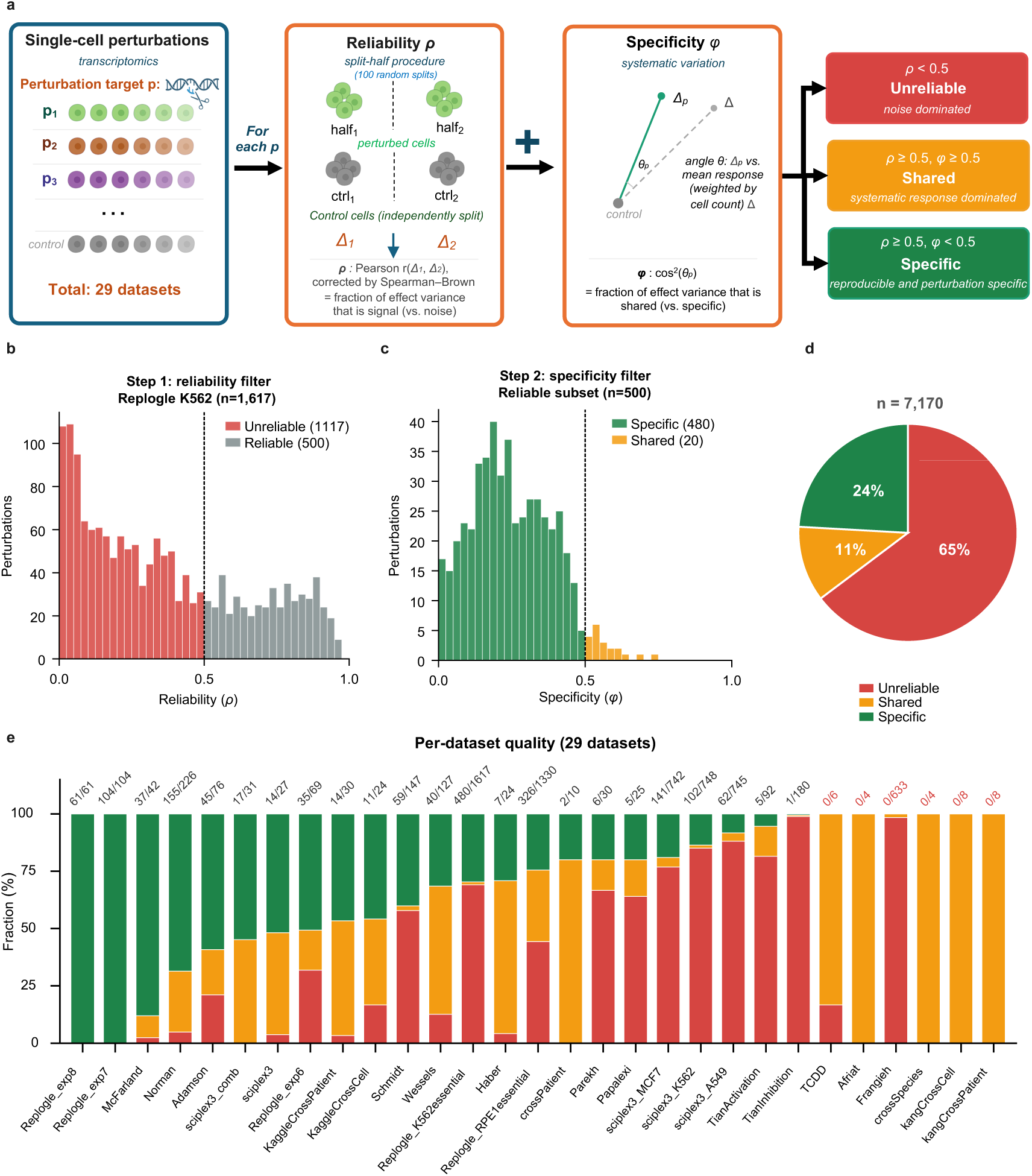
Quality classification for perturbation transcriptomics. **(a)** Workflow: for each perturbation in a Perturb-seq screen, split-half reliability *ρ* is estimated by partitioning perturbed cells into two random halves, computing the per-gene effect vector for each half against an independently split control, and calculating the Pearson correlation between the two vectors across genes. This is repeated over 100 random splits and *ρ* is the Spearman–Brown correction of the median Pearson correlation. Specificity is quantified as *φ* = cos^2^ *θ*, the systematic variance fraction (the squared cosine similarity between the perturbation’s effect vector and the mean perturbed response across cells in all reliable perturbations in the dataset; following ^27^). The two scores are used to classify perturbations as unreliable (*ρ <* 0.5), shared (*ρ* ≥ 0.5 and *φ* ≥ 0.5), or specific (*ρ* ≥ 0.5 and *φ <* 0.5). **(b)** Histogram showing the reliability filter on the Replogle K562 essential-gene dataset ^6^ (number of perturbations *n* = 1,617): histogram of *ρ* across all perturbations; the threshold *ρ* = 0.5 separates 1,117 unreliable from 500 reliable perturbations. **(c)** Histogram showing the second-step specificity filter applied to the 500 reliable perturbation subset: histogram of *φ*; the threshold *φ* = 0.5 separates 20 shared from 480 specific perturbations. **(d)** Pooled summary across all 29 datasets curated from scPerturBench ^26^ (*n* = 7,170 perturbation–condition combinations). **(e)** Per-dataset breakdown across all 29 datasets, sorted by specific fraction. Numbers above each bar give the count of specific perturbations / total perturbations in that dataset.

Across 7,170 perturbations in 29 datasets, 65% were classified as unreliable, 11% as shared, and only 24% as specific (Figure 1d). Several commonly used datasets contained few specific perturbations (Figure 1e). As a robustness check, the scarcity was more pronounced under a reliability measure that compares whole cell distributions rather than pergene averages (Supplementary Figure S1). Stratifying perturbations by cells per condition (Supplementary Figure S2a) showed that increasing the number of cells reduces unreliability sharply (84% unreliable at *<*50 cells; 10% at *>*500 cells) but does not eliminate the shared fraction, which remains around 6–20% at all cell counts.

To further characterize the shared and specific responses biologically, we compared their pathway enrichment profiles (Supplementary Figure S3). The shared response was enriched for a narrow and recurrent set of pathways dominated by hallmark cellular processes such as the cell cycle and proliferation: E2F targets and the G2-M checkpoint were enriched in 11 of the 13 qualifying datasets; Myc targets in 9; and p53, interferon, and mTORC1 signaling in a subset. The specific responses, in contrast, were enriched for a far broader and more numerous set of pathways than the shared response (50 distinct Hallmark programs ^36^ versus 34, and over 1,200 versus 47 Gene Ontology biologicalprocess terms ^37^; Supplementary Figure S3b–e), spanning diverse biology such as hypoxia, hormone (androgen and estrogen) responses, and lipid metabolism rather than a single common program.

### 2.2 Quality filters explain evaluation and improve training efficiency

If most of the evaluation targets are unreliable or shared, existing benchmarks might mostly reflect how well the models predict the mean response across the perturbations. We tested this hypothesis directly by re-analyzing the data from Wei et al. ^26^ and Ahlmann-Eltze et al. ^21^, two recent large-scale perturbation-prediction benchmarks that span eight settings across genetic, cellular, and chemical prediction tasks in single-target or multi-target designs.

When unreliable and shared perturbations were removed from evaluations, the method rankings changed across seven out of eight benchmark configurations, but the magnitude of the change varied (Figure 2a–b, Supplementary Figure S5). The largest reshuffle occurred in the Wei et al. genetic single benchmark, where scGPT ^15^ dropped seven ranks (rank 1 on all perturbations to rank 8 on specific perturbations), GenePert ^38^ rose to rank 1, scELMo ^39^ rose five ranks (rank 7 to 2), and CPA (Compositional Perturbation Autoencoder) ^40^ rose from rank 6 to rank 4. The other seven configurations, including both Ahlmann-Eltze et al. benchmarks, showed smaller shifts, with the top method still changing in four of seven cases. Stratifying evaluation scores by perturbation quality label (Figure 2c) showed a consistent pattern across the four genetic benchmark panels: in 39 of 41 method–benchmark combinations, mean model performance followed Shared *>* Specific *>* Unreliable. Thus, under the published metrics, shared perturbations were generally the highest-scoring category, whereas unreliable perturbations usually yielded the lowest scores. This pattern indicates that aggregate benchmark scores depend on the composition of perturbation quality in the evaluation set.

**Figure 2:**
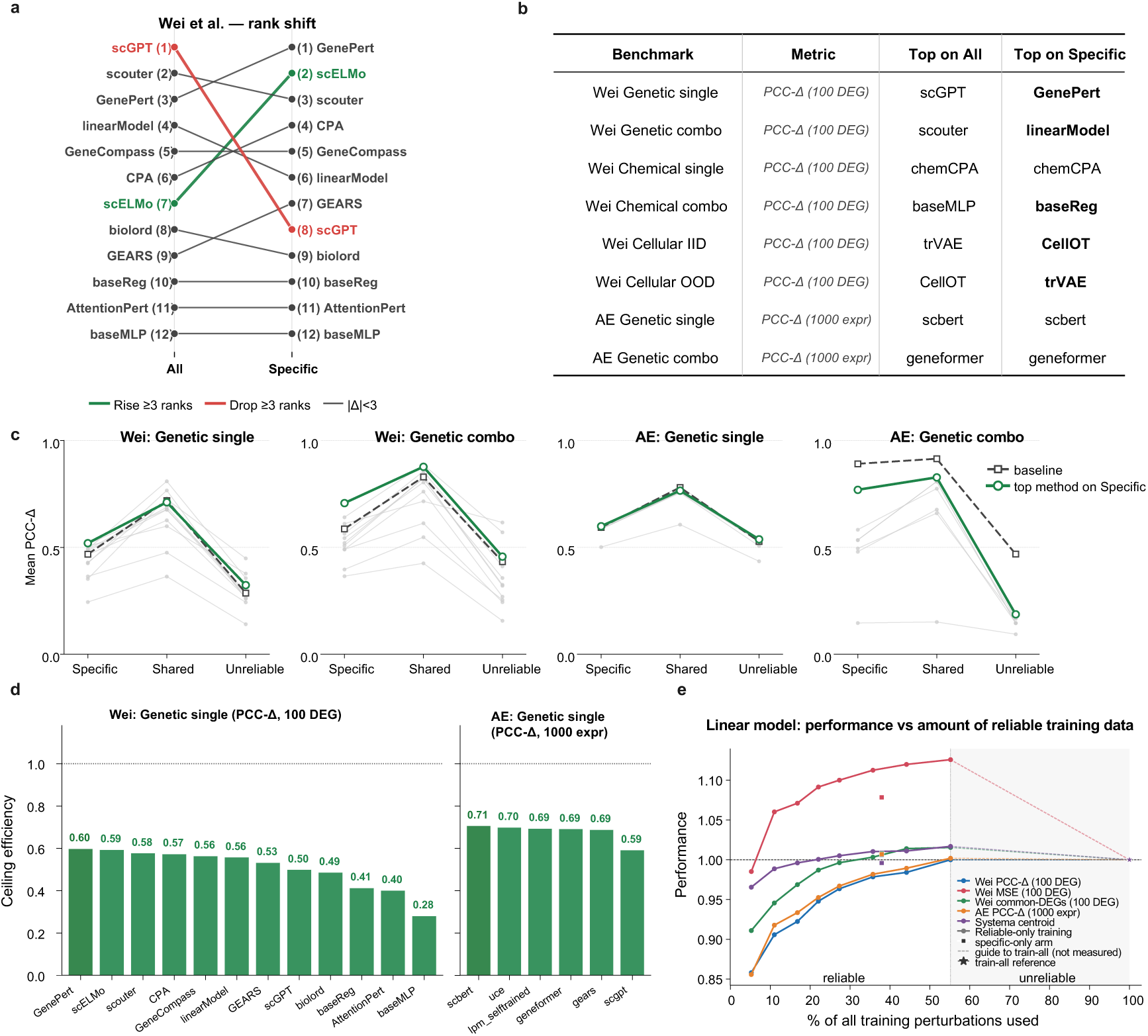
Quality filtering affects benchmarking. **(a)** Method rank changes when restricting evaluation to specific perturbations, for the Wei et al. benchmark. Methods that rise or drop ≥ 3 ranks are highlighted. The scGPT-CPA reversal is consistent across alternative threshold choices (Supplementary Figure S4). **(b)** Top-method change across eight benchmarks from ^21,26^. Pearson correlation was selected as the main metric used in the original studies. While both studies calculated the Pearson correlation between the predicted and the measured change in expression relative to control, Wei et al. ^26^ used the top 100 differentially expressed genes (DEGs) and Ahlmann-Eltze et al. ^21^ used the top 1,000 highly expressing genes in the control condition. For each benchmark, the table reports the top-ranked method on all perturbations vs. on specific perturbations only. Bold font under “Top on Specific” marks benchmarks where the top method changed. **(c)** Per-method dataset-weighted mean performance across perturbation quality classes on four genetic benchmarks in Wei et al. ^26^ and Ahlmann-Eltze et al. ^21^, evaluated on the metrics described in panel (b) caption. The dashed line is each benchmark’s own baseline: the training set mean for the two Wei panels, the Ahlmann-Eltze mean baseline for their genetic single panel, and the additive model for their genetic combo panel. The green line is the top-on-Specific method in each benchmark. **(d)** Ceiling efficiency on specific perturbations (dataset-weighted average). **(e)** Linear-model performance versus the amount of reliable training data, evaluated on held-out specific perturbations (three genetic-single perturbation datasets, 75/25 train/test split, five seeds). The *x*-axis shows the percentage of all training perturbations used; the *y*-axis the performance on the specific test set relative to training on all perturbations (= 1, dashed line). Curves are random reliable subsets swept by size (mean over 10 draws per size; five metrics: Wei PCC-Δ, Wei MSE, Wei common-DEGs ^26^, Ahlmann-Eltze Pearson-Δ^21^, Systema centroid ^27^; Wei MSE plotted as train-all/subset so that up is better for every metric); squares mark the specific-only subset and stars mark training on all perturbations. The shaded band is the unreliable fraction of training data, beyond the full reliable subset.

To separate model error from measurement error in evaluations, we used Spearman’s correction for attenuation ^33,35^. If a perturbation’s measured effect has reliability *ρ*, then even a perfect predictor of the underlying effect can achieve a Pearson correlation with the measured effect of at most 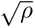. We therefore normalized each observed Pearson correlation *r* by this maximum, defining ceiling efficiency as 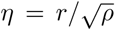. For each method, we first averaged *η* over perturbations within each dataset and then averaged across datasets, so that each dataset contributes equally to the final score. On specific perturbations, ceiling efficiencies clustered within a limited range in both benchmarks: approximately 0.3–0.6 in Wei et al. ^26^ and 0.6–0.7 in Ahlmann-Eltze et al. ^21^ (Figure 2d). Thus, after restricting the evaluation to specific targets and normalizing by their measurement reliability, no method separated strongly from the others, reinforcing recent reports that complex architectures do not substantially outperform simple linear baselines on perturbation prediction ^20–22,24,31,32^.

We then asked whether the framework also could provide an immediate, actionable improvement to current training pipelines. We re-trained the linear model from Wei et al. ^26^ on nested subsets of the training perturbations — all perturbations, the reliable subset, and the specific subset — under a 75/25 train/test split repeated across five seeds in three genetic datasets, and evaluated every arm on the same held-out specific test perturbations (Figure 2e). Across all five metrics, training on the reliable subset (∼55% of all perturbations) matched or exceeded training on all perturbations, including a ∼13% improvement in Wei MSE ^26^. Quality labels therefore identify higher-value training targets: most of the predictive signal for specific perturbations is carried by the reliable and specific subsets, and the unreliable majority can be removed with little effect. The same conclusions broadly held as the training set was scaled (Supplementary Figure S6) and across complex models (Supplementary Figure S7).

Together, these results show that perturbation-level quality labels are useful for both evaluation and training. For evaluation, the labels separate prediction of broad shared responses from prediction of reliable and specific effects, so benchmark scores can be interpreted in relation to the target class being tested. For training, quality labels identify the higher-value targets: the reliable and specific subsets recover full-data performance on specific perturbations with substantially fewer training examples, so reliability filtering improves the data efficiency of training. Furthermore, as unreliable training perturbations contribute little to model performance (Figure 2e) and performance increases with more reliable training data (Supplementary Figure S6), improving perturbation prediction models requires generating more reliable perturbations, not simply more data.

### 2.3 A one-parameter reliability model enables prospective experimental sample-size planning

By the law of large numbers, perturbations with more cells tend to be more reliable (Supplementary Figure S2), indicating that sampling depth can improve the reliability of pseudo-bulk effect estimates. The practical question for experimentalists therefore becomes how many cells per perturbation are needed to reach the reliability threshold *ρ* ≥ 0.5. In Perturb-seq experiments where cost scales with the number of cells sequenced, the answer matters directly for screen design.

Under the standard assumption that each cell contributes an independent measurement of the perturbation effect, classical test theory predicts how reliability scales with cell count ^33–35,41^. If *τ* ^2^ denotes the per-cell signal-to-noise ratio for a perturbation, then reliability increases with the number of cells *N* as

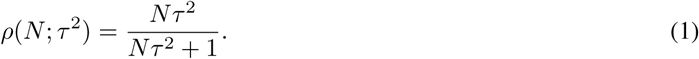

This simple one-parameter model captured empirical reliability curves with high fidelity (Figure 3a–c). Across 5,343 perturbations with at least 100 cells, the median fit of this model to each perturbation’s own reliability curve was *R*^2^ = 0.95 (Figure 3c; per-dataset fits in Supplementary Figure S8). The fitted *τ* ^2^ values varied widely across perturbations and datasets, yielding large differences in the cell count required to achieve *ρ* ≥ 0.5.

**Figure 3:**
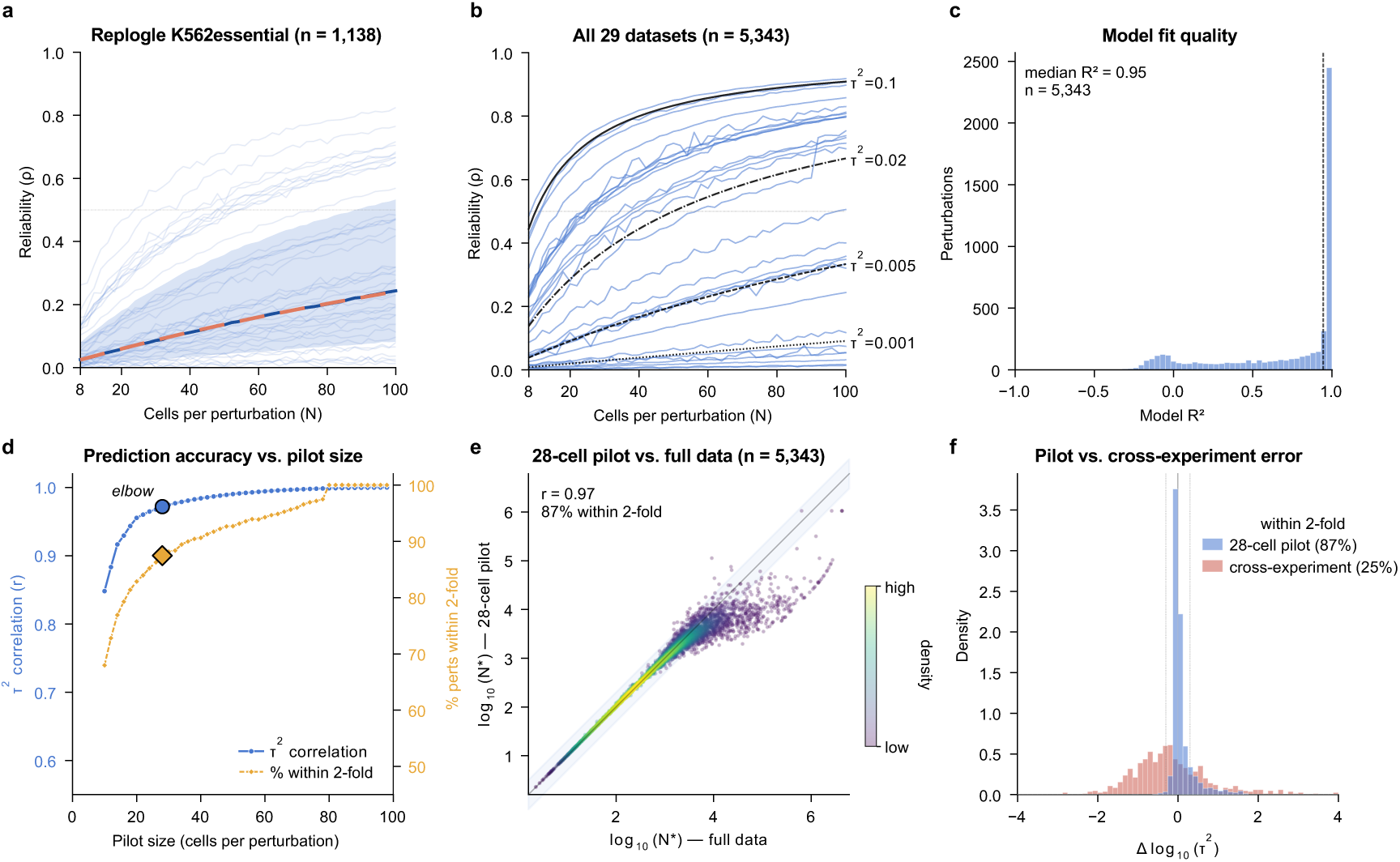
Experimental screen design based on a model for reliability as a function of sample size. **(a)** Per-perturbation reliability curves for Replogle K562 essential-gene dataset (*n* = 1,138 perturbations with at least 100 cells, each fit separately with its own *τ* ^2^). Faint blue lines: individual perturbations’ empirical curves; solid dark blue line: per-*N* median across perturbations. The dashed orange line shows the parametric form *ρ*(*N*) = *Nτ* ^2^*/*(*Nτ* ^2^ + 1) evaluated at the median of the 1,138 fitted *τ* ^2^ values, indicating a single representative curve for visualization rather than a global fit. **(b)** Per-dataset median reliability curves for all 29 datasets overlaid on parametric reference curves at *τ* ^2^ 0.001, 0.005, 0.02, 0.1 . **(c)** Validation of the parametric functional form: distribution of *R*^2^ between each perturbation’s empirical reliability curve and its individual parametric fit (median *R*^2^ = 0.95 across *n* = 5,343 perturbations). **(d)** Pilot accuracy versus size: correlation between pilot *τ* ^2^ and full-data *τ* ^2^ (blue) and fraction of *N ^∗^* predictions within two-fold of the full-data estimate (orange). The elbow falls around 28 cells per perturbation. **(e)** Pilot-predicted *N ^∗^* vs. full-data *N ^∗^* at the 28-cell pilot (*r* = 0.97, 87% within two-fold; *n* = 5,343). **(f)** Pilot prediction error vs. cross-experiment transfer error (28-cell pilot vs. borrowing *τ* ^2^ from a different dataset profiling the same perturbation).

We then asked whether *τ* ^2^ can be estimated from a small pilot experiment, as this would inform the optimal design of the full screen. Across the same *n* = 5,343 perturbations, the elbow of pilot accuracy occurred around 28 cells per perturbation (Figure 3d). At this pilot size, the required cell count *N ^∗^* (the minimum number of cells needed to reach *ρ* ≥ 0.5) was predicted within two-fold of the full-data estimate for 87% of perturbations (Figure 3d–e), with the remaining failures concentrated among perturbations with very large *N ^∗^*. Pilot sizes larger than 28 cells produced diminishing returns, with a 60-cell pilot reaching 94% within two-fold versus 87% at 28 cells (Figure 3d).

Pilot-based planning also outperformed cross-experiment transfer. Borrowing *τ* ^2^ from a different dataset profiling the same perturbation kept only 25% of perturbations within two-fold accuracy, compared to 87% for a 28-cell pilot (Figure 3f), despite moderate cross-experiment agreement (intraclass correlation coefficient ≈ 0.69). Even a minimal same-screen pilot therefore provides sample-size guidance that cannot be reliably obtained from published screens alone.

## 3 Discussion

Perturbation prediction has become a flagship task for virtual cell efforts, but recent benchmarking studies have reached pessimistic and sometimes conflicting conclusions about whether deep learning and foundation models improve over simple baselines. Our results show that these disagreements largely trace back to the assumption of those studies that the measured perturbation effect is reliable and perturbation-specific. Across 29 widely used datasets, perturbation-specific signal is present in only a minority of perturbations (24%). Once benchmarks are restricted to specific perturbations, method rankings reshuffle, effect sizes shrink, and the remaining gap to perfect prediction is better interpreted through a reliability-imposed ceiling.

These findings have direct implications for benchmarking practice. Reporting a single score averaged over all perturbations is often misleading because unreliable, shared, and specific perturbations reflect different aspects of model performance. As a minimal standard, we suggest (*i*) reporting the fraction of specific perturbations in the evaluation set, (*ii*) stratifying the results by quality category, and (*iii*) reporting ceiling-normalized metrics (e.g., ceiling efficiency) on the specific subset. A benchmark of perturbation prediction is intended to measure the perturbation-specific response, yet shared perturbations reward prediction of the average response across perturbations, and unreliable ones have no stable signal to predict. Only the specific subset reflects the intended quantity. Community challenges such as the Virtual Cell Challenge ^2^ could therefore adopt reliability-based quality filtering as a default preprocessing step.

For experimental design under a fixed budget, allocating additional cells to fewer perturbations can convert unreliable targets into reliable ones. A 28-cell experimental pilot is sufficient to predict the required sample size of a reliable full-scale experiment for most perturbations.

Recent work showing that calibrated DEG-aware metrics can restore an advantage for deep learning models ^29,30^ addresses the complementary problem of metric design. Our framework operates upstream by asking whether the evaluation target itself is informative. In principle, both approaches can be combined: reliability filtering selects perturbations with stable ground truth, while calibrated metrics ensure that model comparisons on those perturbations reflect biologically relevant differences. Concurrent work has used per-cell-line split-half experimental reliability as a performance reference for prediction models ^42^. Our framework complements this by providing per-perturbation reliability and applying the Spearman–Brown correction for attenuation to convert it into a theoretical ceiling, identifying which targets are reliable enough to serve as ground truth in the first place, and supplying a design rule for generating more reliable targets prospectively.

There are several limitations in our study. Our reliability score measures whether a perturbation’s average effect is stable when its cells are resampled. A low score therefore does not mean the perturbation is biologically uninformative; instead it means that, given the cells sequenced, the average effect cannot be estimated reliably enough to serve as a target for training or evaluation. Neither score reflects biological validity. A confounding effect shared by all cells of a single perturbation, such as a batch or capture effect, can raise the reliability score and still pass the specificity filter, so neither score on its own distinguishes a true perturbation effect from a confounding factor. Our specificity score *φ* follows Viñas Torné et al. ^27^ and captures an important shared component, but other confounders, such as unmodeled covariates or compositional shifts, can bias evaluation in the same way. More broadly, evaluations that do not account for these experimental sources of variation will conflate confounding with model performance, making benchmark conclusions unreliable and even misleading regardless of model complexity. The thresholds we use (*ρ* = 0.5, a 1:1 ratio of reliable signal to noise, and *φ* = 0.5, a 1:1 ratio of perturbation-specific to shared variance) are principled and interpretable, and our main findings are robust above the reliability threshold (*ρ* ≥ 0.5), though below it the criterion admits noise-dominated targets and rankings can shift (Supplementary Figure S4). Specific applications may warrant different operating points depending on the target reliability and the experimental or modeling context.

Looking forward, the low fraction of specific perturbations across existing datasets points to improvements on three fronts. Experimentally, screens could prioritize measurement depth over breadth, allocating more cells per perturbation to raise the fraction that reach a reliable signal. Computationally, models and metrics could use the two quality estimates directly: weighting each perturbation by its reliability so that noisy targets contribute less, and representing the shared and perturbation-specific responses as separate components so that models are evaluated on perturbation-specific biology rather than credited for predicting the shared response. For benchmarking practice, treating data quality as an explicit variable in benchmark construction, rather than an implicit assumption, may be necessary to separate genuine model improvements from differences that arise only because the evaluation targets are unreliable. Beyond perturbation prediction in single-cell transcriptomics, the issues identified here reflect a broader challenge in applying machine learning to biology. Benchmark targets are often derived from noisy biological measurements, which place a ceiling on the accuracy any model can reach, so a model’s remaining error against such a target reflects both the limits of the measurement and the limits of the model. The split-half reliability framework developed here is, in principle, applicable to other settings where measurement profiles serve as ground truth.

In summary, we provide a reliability framework that makes perturbation ground-truth quality explicit and quantitative, explains why many published model leaderboards are unstable, improves model training efficiency, and supplies a simple experimental-design rule that is feasible in practice.

## Methods

### Datasets

We analyzed 29 preprocessed single-cell perturbation datasets from scPerturBench (Wei et al. ^26^). Data acquisition and preprocessing procedures are described in the original publication. We treated each perturbation in our analysis as a perturbation–condition combination, defined as a unique pairing of a perturbing agent or combination of agents with the condition in which the response was measured, comprising the biological context (for example, cell line, cell type, patient, or species) and, where applicable, the experimental context (for example, dose or time point).

### Split-half reliability

For each perturbation with *N* perturbed cells, we estimated the reliability of the pseudo-bulk effect vector ***δ*** ∈ R*^G^*, the change in mean expression of each gene between the perturbed and control cells, using a split-half procedure. Writing the observed effect for gene *g* as *δ_g_*= *δ*_true*,g*_ + *ε_g_*, with measurement error *ε_g_* uncorrelated with the true effect, reliability is the share of the observed variance across genes that comes from the true effect, *ρ* = Var[*δ*_true_]*/*Var[*δ*] = 1−Var[*ε*]*/*Var[*δ*]. Perturbed cells were randomly partitioned into two halves of size ⌊*N/*2⌋, and the perturbation effect in each half was computed as the difference in mean expression relative to a similarly split control population. The Pearson correlation *r*_half_ between the two half-sample effect vectors was computed over *K* = 100 random partitions. Its median across partitions is denoted by *r̃*_half_ . The Spearman–Brown prophecy formula ^34,43^ relates reliability *ρ* to this median Pearson correlation between split-halves as

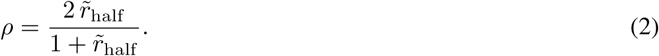

Perturbations with *ρ* ≥ 0.5 were classified as reliable. Reliability was computed at fixed subsample sizes *n*_half_ ∈ {8, 16, 32*, . . .,* 512} and at the maximum available cell count, and additionally on a denser grid *n*_half_ ∈ {4, 5, 6*, . . .,* 50}, or equivalently *n*_total_ = 2*n*_half_ from 8 to 100 cells, for parametric fitting.

As a robustness check, we also computed split-half reliability using the energy distance, a non-parametric multivariate distance between two empirical distributions ^44^, established as a per-perturbation metric for single-cell perturbation data ^28^. For two sets of cells *X* = {*x*_1_*, . . ., x_n_*} and *Y* = {*y*_1_*, . . ., y_m_*} in gene-expression space,

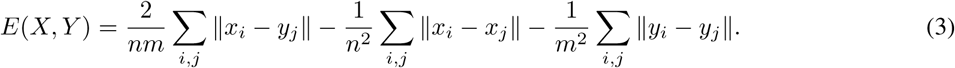

For each perturbation, we defined a distributional analog of reliability

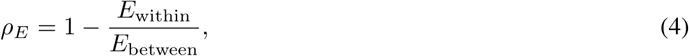

where *E*_within_ is the median energy distance between two random equal-sized halves of the perturbed cells across *K* = 100 resampling seeds, and *E*_between_ is the energy distance between the full perturbed and control cell pools. We applied this to the same 29 datasets, using *ρ_E_* ≥ 0 as the cutoff for reliability, the threshold where the perturbation effect at least equals within-perturbation dispersion. Specificity for this robustness check was kept the same.

### Quality classification

Each perturbation was classified by combining reliability with systematic variation. We quantified systematic variation as the cosine similarity cos *θ* between the perturbation’s effect vector and the mean effect across all reliable perturbations in the same dataset, following ^27^. We then defined the systematic variance fraction *φ* = cos^2^ *θ* ∈ [0, 1], the squared cosine similarity, which equals the fraction of the perturbation effect vector’s variance projected onto the systematic axis (*φ* ≈ 1 indicates a perturbation whose effect is dominated by the systematic axis; *φ* ≈ 0 indicates a perturbation-specific effect orthogonal to it). Both classification thresholds were chosen such that each corresponds to a 1:1 variance ratio. The reliability threshold *ρ* = 0.5 follows the convention from classical test theory ^33,35^: half of the variance in the pseudo-bulk profile is signal, half is measurement noise. The specificity threshold *φ* = 0.5 corresponds to a 1:1 ratio of perturbation-specific to systematic variance: half of the perturbation effect vector’s variance is shared with other perturbations in the dataset, half is perturbation-specific. Unreliability was assessed first: perturbations with *ρ <* 0.5 were classified as unreliable regardless of their systematic-variation score. Among the remaining reliable perturbations (*ρ* ≥ 0.5), those with *φ* ≥ 0.5 were classified as shared, and the remainder as specific.

### Benchmark re-evaluation

We obtained published per-perturbation benchmark results from Wei et al. ^26^ and Ahlmann-Eltze et al. ^21^ . The Wei et al. benchmark spans six configurations: genetic single (9 datasets, 15 methods), genetic combo (4 datasets, 15 methods), chemical single (3 datasets, 9 methods), chemical combo (1 dataset, 6 methods), cellular IID (12 datasets, 14 methods) and cellular OOD (12 datasets, 14 methods); all are scored with Pearson correlation on the top 100 differentially expressed genes per perturbation. The Ahlmann-Eltze et al. benchmark spans two configurations: genetic single (3 datasets, 7 methods in the main evaluation and 16 in the architecture ablation) and genetic combo (1 dataset, 9 methods); both are scored with the same metric, the Pearson correlation of the effect (delta) vector (the authors’ “Pearson delta”) of the top 1000 highly expressed genes in the control condition. In both cases, we used the authors’ published result tables directly, merging per-perturbation scores with our quality labels. Perturbations were stratified by quality category and method rankings were recomputed within each stratum by their mean excess over that benchmark’s baseline, averaged across datasets. Ranking exclude each benchmark’s baseline.

### Ceiling efficiency

Under Spearman’s correction for attenuation ^33,35^, the observed Pearson correlation between two error-prone measurements *X*_obs_ and *Y*_obs_ with reliabilities *ρ_X_*and *ρ_Y_* relates to the correlation between their underlying true scores as

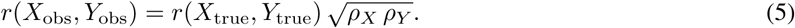

In our setting, *X* is the deterministic output of a trained model (so *ρ_X_* = 1) and *Y*_obs_ is a noisy pseudo-bulk measurement whose reliability *ρ* we estimate by split-half resampling. The formula then reduces to 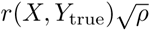. Since |*r*(*X, Y*_true_)| ≤ 1, the observable correlation is bounded by 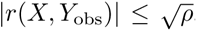, with equality when *X* is a perfect estimate of *Y*_true_. For a correlation-based metric we therefore defined ceiling efficiency as 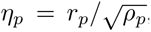, where *r_p_* is the Pearson correlation between model prediction and ground truth for perturbation *p*. Because Ahlmann-Eltze et al. also score predictions with a Pearson correlation of the delta effect vector, the same bound applies and we used the identical normalization 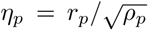 for both benchmarks. In both cases *η* = 1 indicates performance at the theoretical ceiling imposed by measurement noise. Ceiling efficiencies were computed on specific evaluation perturbations only. Per-method ceiling efficiencies reported in the main text and Figure 2d are dataset-weighted averages. For each method, we computed the mean *η_p_* within each dataset, then averaged across datasets, so each dataset contributes equally regardless of its perturbation count.

### Retraining on quality-filtered subsets

To test whether the perturbation quality labels identify higher-value training data, we re-trained the linear model of Wei et al. ^26^ on nested subsets of the training perturbations and evaluated every subset on a common held-out test set. We restricted this analysis to three genetic single Perturb-seq datasets (Replogle K562-essential, Replogle RPE1- essential, and Adamson). These were selected because they are genetic perturbation screens that contain all three quality classes — so that the training arms differ — and at least 15 specific perturbations, enough to form a held-out specific test set; datasets that are chemical, contain no specific perturbations, or are composed almost entirely of a single class were excluded from the retraining analysis but remain in the reliability characterization (Figure 1). For each dataset we split perturbations 75/25 into training and test sets, stratified by quality label and perturbation type, and repeated the split across five random seeds. The held-out test set was identical across all training arms within a split, and no test perturbation appeared in any training set, so the model never saw a test perturbation during training. We defined training arms nested by quality: all (every training perturbation), reliable (reliable perturbations only), and specific (specific perturbations only). For the curve in Figure 2e, we additionally trained on random subsets of the reliable pool drawn across a range of sizes (ten draws per size). Each trained model predicted the same held-out test perturbations, and we aggregated per-perturbation scores over the specific test perturbations, the primary evaluation target (Figure 2e. The reliable and specific test pools are compared in Supplementary Figure S6). We scored each test perturbation with five metrics: the Pearson correlation of the top-100 differentially expressed genes’ effect vector (Wei PCC-Δ) ^26^, the mean squared error over those genes (Wei MSE) ^26^, the number of shared differentially expressed genes (Wei common-DEGs) ^26^, the Ahlmann-Eltze delta Pearson correlation over the top-1000 expressed genes ^21^, and the Systema cosine-weighted centroid accuracy ^27^. In Figure 2e, performance is expressed relative to training on all perturbations (= 1), with Wei MSE plotted as the train-all/subset ratio so that higher is better for every metric. To test whether the same ordering holds for higher-capacity models, we repeated the all, reliable, and specific retraining for GEARS ^12^ and scGPT ^15^ and report the training-arm × test-pool grid in Supplementary Figure S7. For these deep models, a small validation slice (about 15% of the specific training perturbations) was held out of the training portion for early stopping, identical across arms and disjoint from the test set, whereas the linear model used no validation set. To characterize how performance scales with the amount of training data, we additionally swept each training pool (reliable, specific, all, and unreliable) across a range of subset sizes and report the resulting curves on the reliable and specific test pools in Supplementary Figure S6.

### Parametric reliability model

Under a random effects model in which the true effects across genes have variance 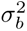 and the cell-to-cell noise has variance 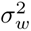, averaging *N* cells leaves 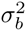 unchanged and reduces the noise variance to 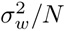 */N*, so the reliability of a pseudo-bulk estimate from *N* cells is 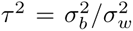, where 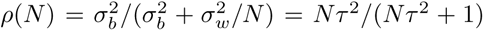. This is the Spearman–Brown formula applied to the mean of *N* measurements ^34,41^. We estimated *τ* ^2^ for each perturbation by nonlinear least-squares fitting of the observed (*N, ρ*) pairs from the fixed-subsample reliability curves (minimum 2 distinct *N* values per perturbation), restricted to perturbations with at least 100 perturbed cells in the source experiment. The required sample size for target reliability *ρ*_0_ is then *N ^∗^* = *ρ*_0_*/*[*τ* ^2^(1 − *ρ*_0_)].

### Pilot validation

To simulate prospective experimental design, we fit *τ* ^2^ using only reliability data up to a pilot budget (10 to 98 cells per perturbation in steps of 2) and compared to the full-data *τ* ^2^. Pilot fitting used the same least-squares procedure with a minimum of 2 data points, with the constraint that at least one observed data point beyond the pilot budget existed for leave-future-out validation. The pilot size at which prediction accuracy plateaus was identified as the elbow of the *τ* ^2^- correlation curve, which fell at 28 cells. The within-2-fold curve elbow agreed closely. The kneedle elbow is defined by rescaling *x* (pilot size) and *y* (accuracy) to [0, 1] and choosing the point that maximizes *y* − *x*, i.e. the point on the accuracy curve farthest from the straight line joining its endpoints, which identifies the inflection where marginal accuracy gains begin to level off. We measured agreement by the Pearson correlation of log_10_(*τ* ^2^) values and the fraction of *N ^∗^* predictions falling within two-fold of the full-data estimate. For genes appearing in multiple datasets, we also compared pilot errors to cross-experiment transfer errors and quantified cross-experiment consistency using the intraclass correlation coefficient of log_10_(*τ* ^2^).

## Code and Data Availability

All scripts used for the analysis of the paper are available at https://github.com/cbg-ethz/scReliability_ paper.git. All datasets used in this study are publicly available from published sources.

## Acknowledgments

This work was supported by Dr. Walter and Edith Fischli via the ETH Zurich Foundation (Project No. 2024-HS-254), and was made possible in part by grant numbers CZIF2024-010302 and CZIF2025-011050 from the Chan Zuckerberg Initiative Foundation.

## Supplementary Materials

- **Supplementary Figure S1:** Distributional reliability robustness check: pooled and per-dataset triage composition under split-half energy-distance reliability.
- **Supplementary Figure S2:** Quality fractions by cells per condition (panel a, pooled across 29 datasets) and per-dataset reliability distributions (panel b).
- **Supplementary Figure S3:** Shared versus specific responses of reliable perturbations: shared-axis Hallmark program enrichment, and the number and breadth of specific-residual programs (Hallmark and GO-BP).
- **Supplementary Figure S4:** Threshold robustness of quality classification (rank differences and specific fraction across the *ρ* × *φ* threshold grid).
- **Supplementary Figure S5:** Method rank change (All → Specific) across the seven benchmark settings other than Wei genetic single (which is shown in Figure 2a).
- **Supplementary Figure S6:** Data-scaling by training pool, evaluated on reliable and specific test perturbations.
- **Supplementary Figure S7:** Multi-model retraining grid: mean performance for each training arm × test pool across five metrics and three models.
- **Supplementary Figure S8:** Per-dataset parametric reliability fits across all 29 datasets.

---

**Figure S1:**
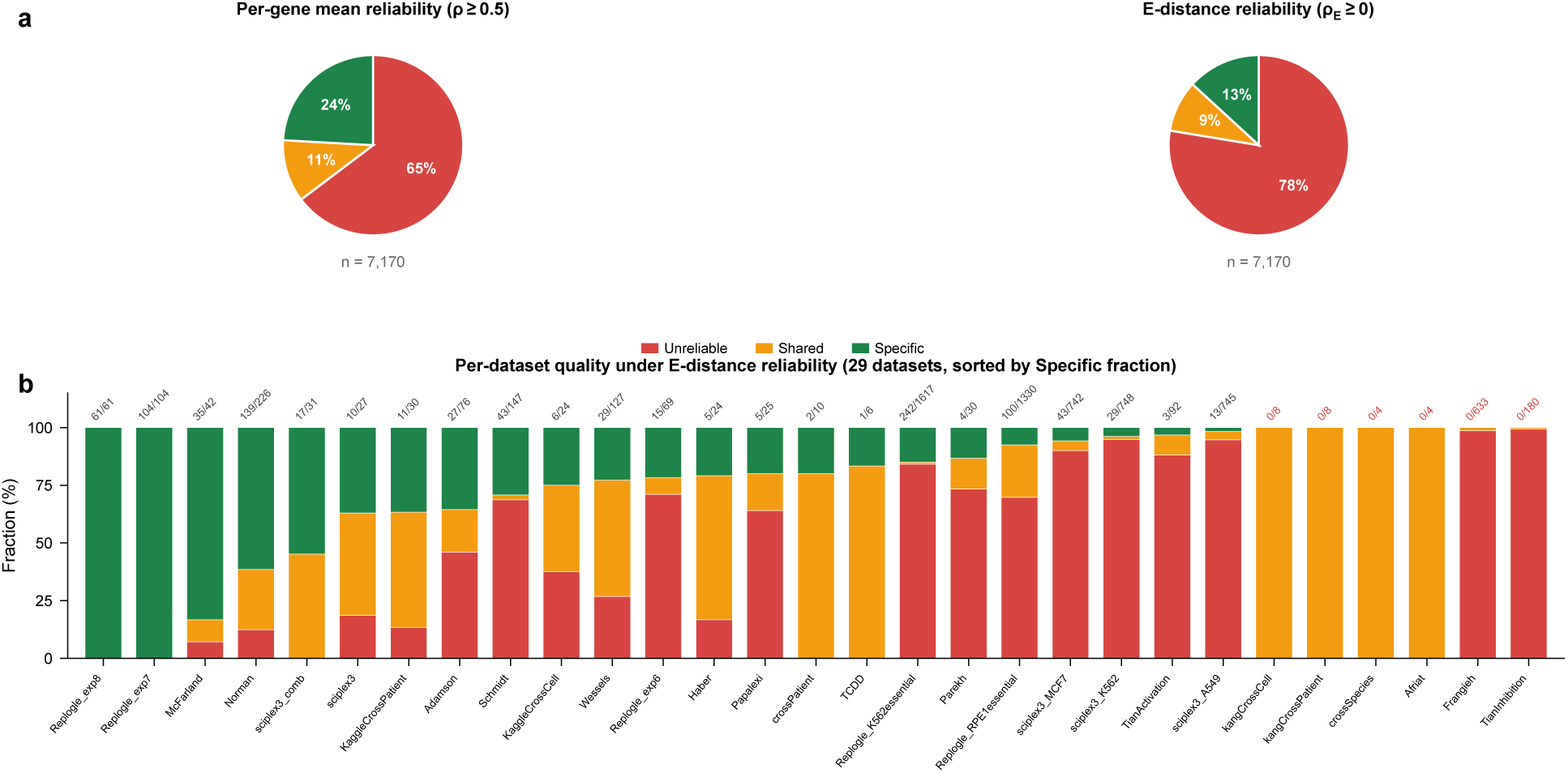
The scarcity of reliable, perturbation-specific data is more pronounced, not less, under a distributional reliability measure. **(a)** Pooled triage composition across all 7,170 perturbation–condition combinations in 29 datasets, computed under two reliability statistics: per-gene mean reliability (Pearson split-half, *ρ* ≥ 0.5; main-text framework, left) and distributional reliability (energy-distance split-half on cell distributions, *ρ_E_* ≥ 0; right). E-distance reliability is more conservative because it penalizes within-perturbation cell-state variance that the per-gene mean averages out. The qualitative conclusion (perturbation-specific data are the minority) holds under either choice and is in fact strengthened under the distributional measure. **(b)** Per-dataset composition under E-distance reliability, sorted by specific fraction. The same datasets that were dominated by unreliable perturbations in the main-text per-gene-mean analysis (Figure 1e) remain dominated by unreliable perturbations here, and the specific fraction is smaller or equal in 28 of 29 datasets with the exception of TCDD. Numbers above each bar give the count of specific perturbations / total in that dataset under E-distance triage.

**Figure S2:**
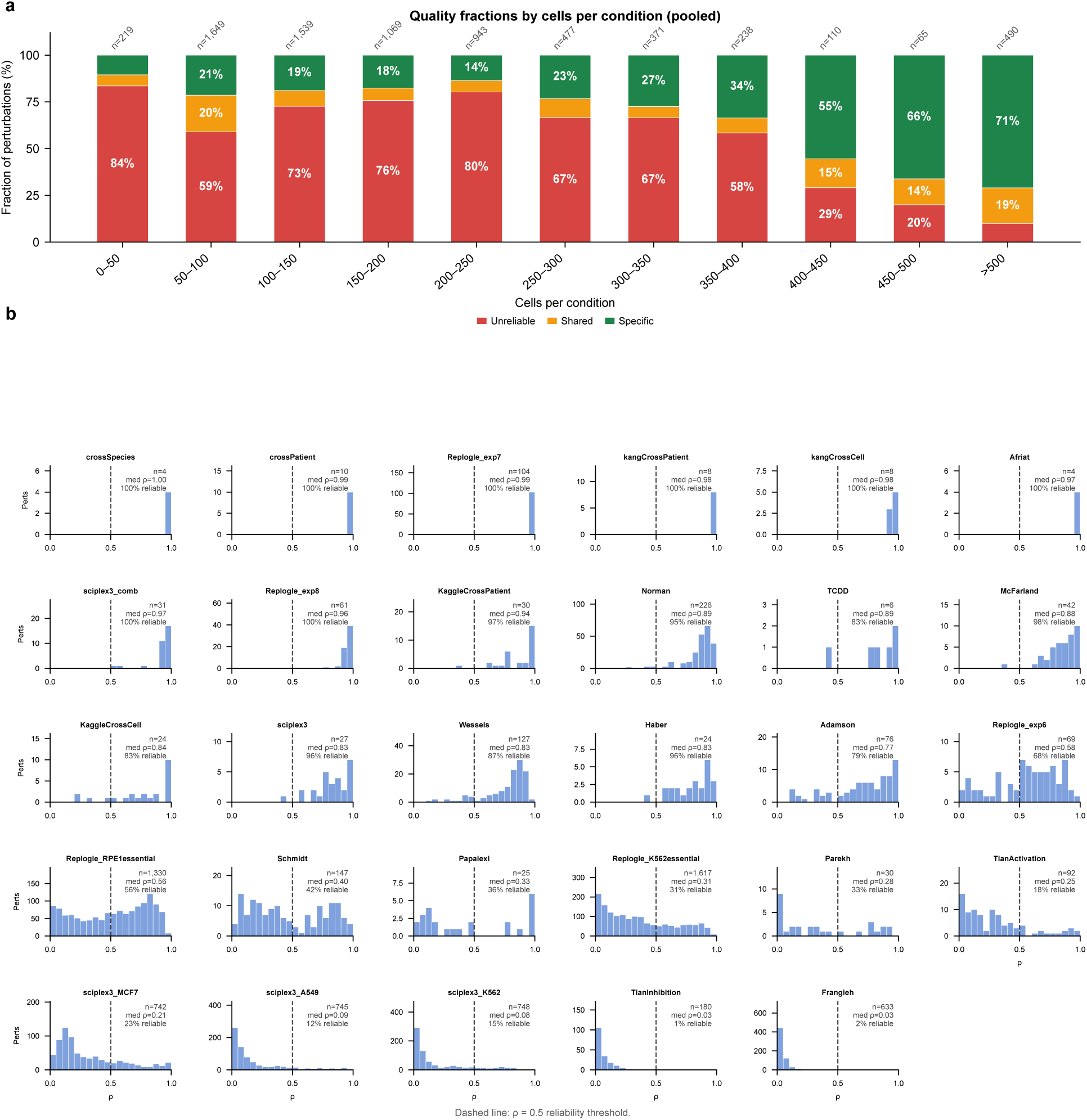
Quality fractions by cells per condition and per-dataset reliability distributions. **(a)** Triage fractions as a function of cells per condition (50-cell bins from 0 to 500, plus a *>*500 bin; pooled across all 29 datasets, genetic and cellular). Sample size shifts the distribution of categories but does not eliminate the shared fraction at any cell count: more cells reduce unreliability sharply but the shared fraction remains in a 6–20% band. **(b)** Histograms of split-half reliability *ρ* for each of the 29 datasets, sorted by median *ρ*, with the threshold *ρ* = 0.5 marked. Genetic datasets in blue, cellular-context datasets in red.

**Figure S3:**
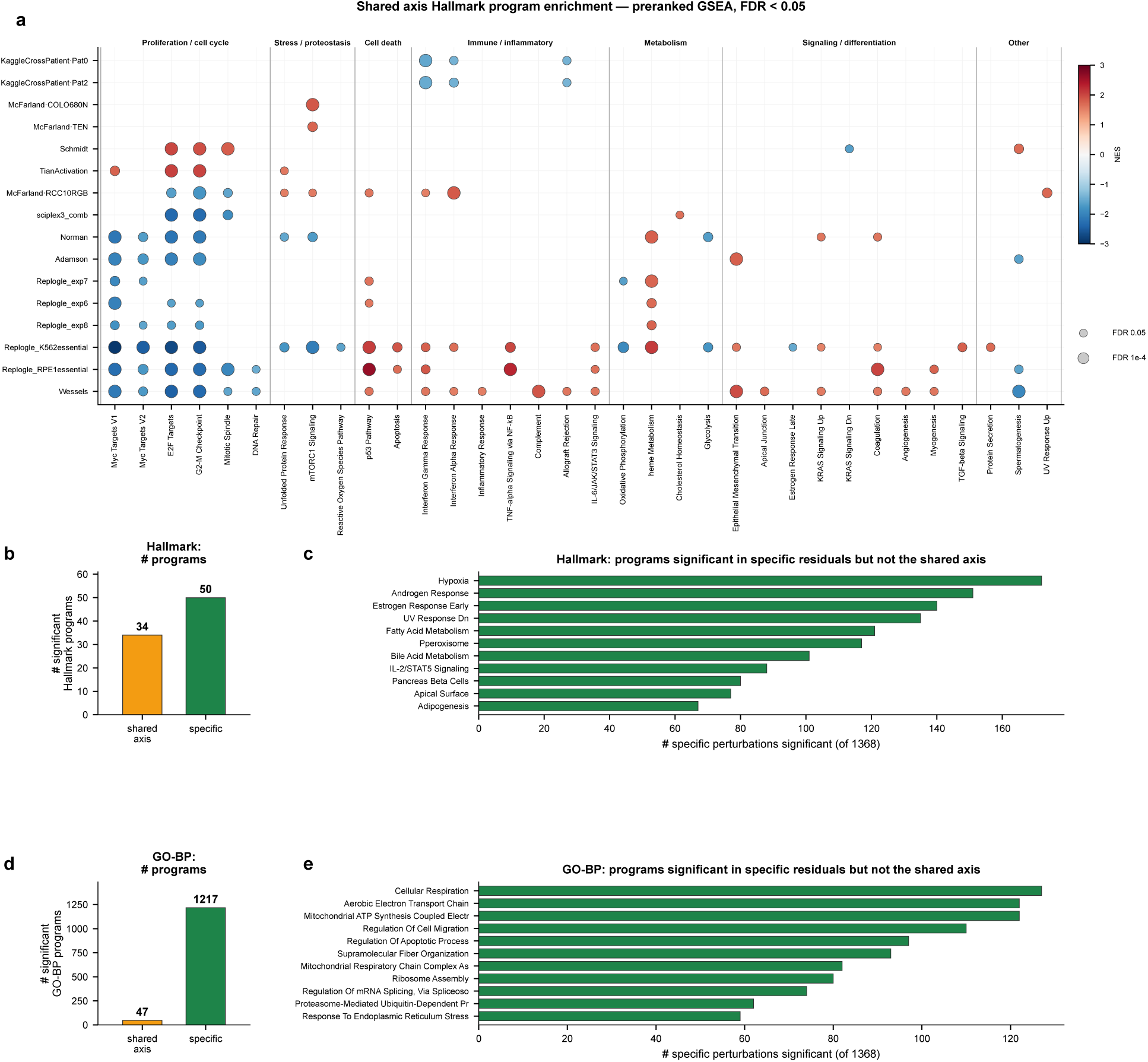
Shared versus specific responses of reliable perturbations. **(a)** Preranked GSEA of the shared-axis gene loadings against Hallmark programs, for each qualifying (dataset, context) with at least 10 reliable perturbations and a coherent shared axis (at least one Hallmark program at FDR *<* 0.05). Rows are ordered by enrichment profile (hierarchical clustering), columns are grouped by program category; dot color is the normalized enrichment score (NES) and dot size is log_10_ FDR. **(b,d)** Number of distinct programs significant (FDR *<* 0.05) along the shared axis versus in the specific residuals, for Hallmark (b) and GO Biological Process (d). **(c,e)** Programs significant in the specific residuals but not the shared axis, ranked by the number of perturbations, for Hallmark (c) and GO-BP (e). The shared axis is dominated by a small set of proliferation/stress programs, whereas the specific residuals spread across a broader and more numerous set of programs.

**Figure S4:**
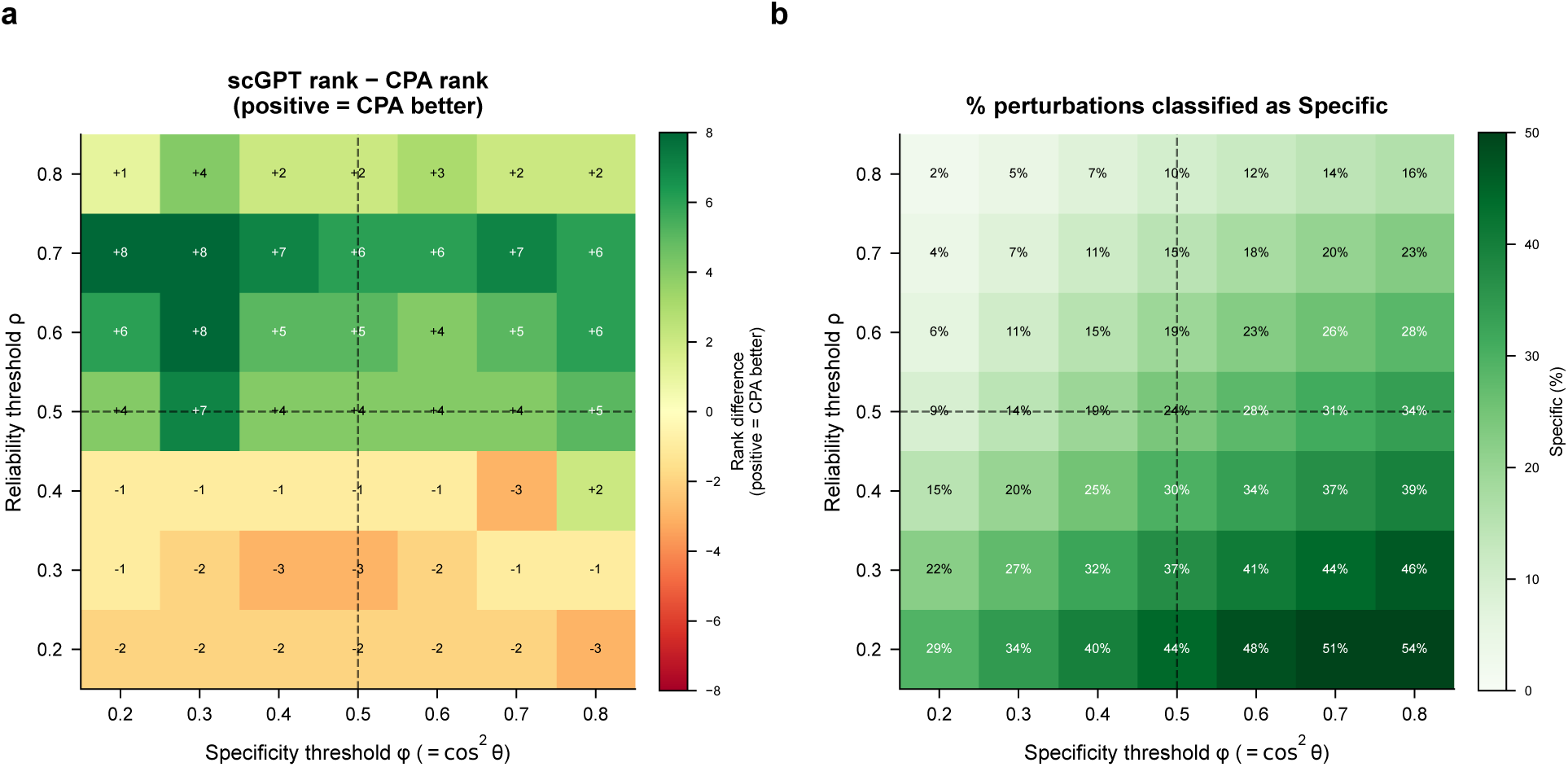
Threshold robustness of quality classification. **(a)** Rank difference between scGPT and CPA on specific perturbations across all combinations of reliability threshold (*ρ*, *y*-axis) and specificity threshold (*φ*, *x*-axis). Positive values (green) indicate that CPA outranks scGPT; the finding holds consistently across the entire region *ρ* ≥ 0.5. **(b)** Fraction of perturbations classified as specific at each threshold pair. Dashed lines mark the thresholds used in the main text (*ρ* = 0.5 and *φ* = 0.5)

**Figure S5.**
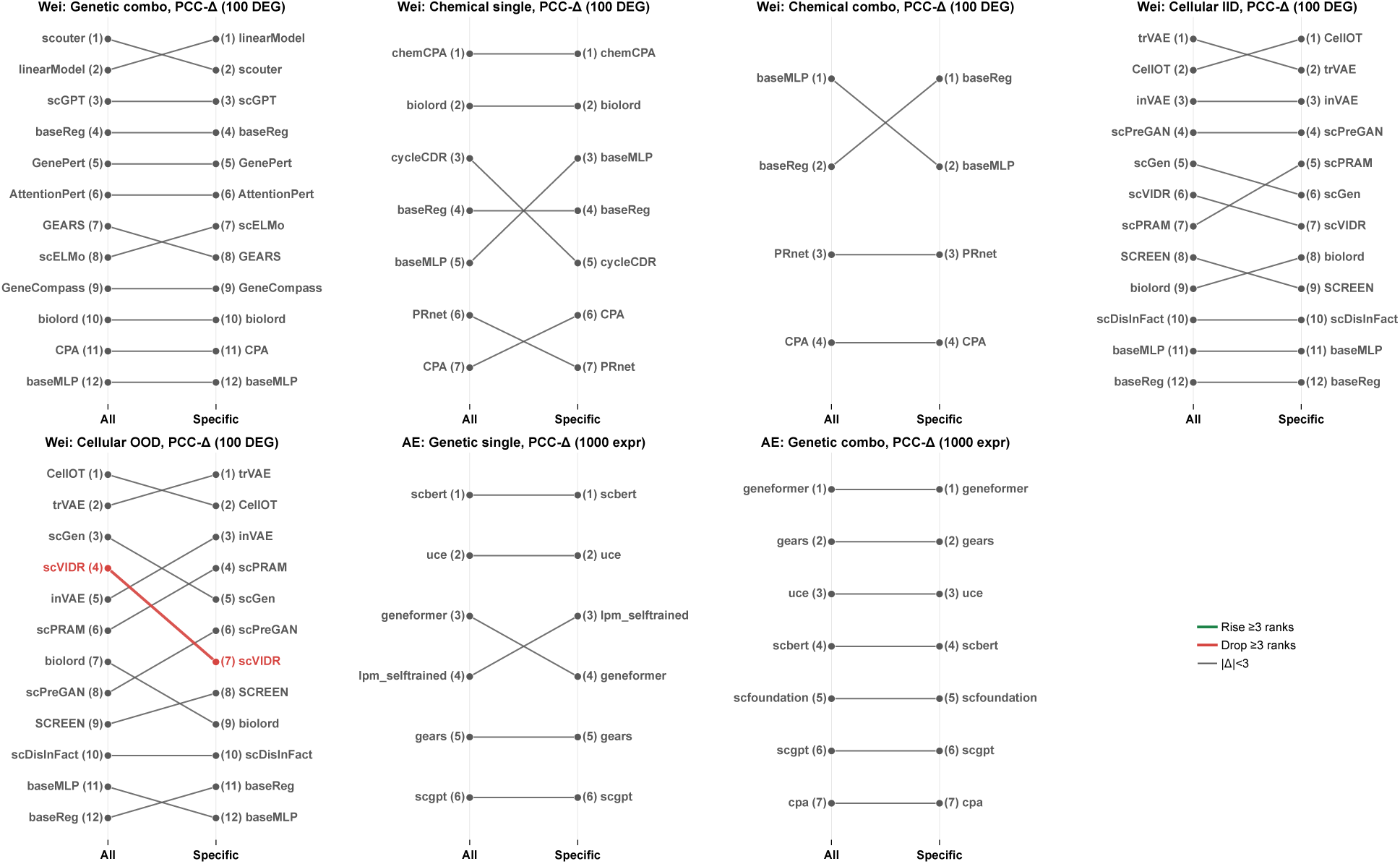
: **Method rank change (All → Specific evaluation perturbations) across the seven benchmark settings other than Wei genetic single.** Wei genetic single, the largest reshuffle (a seven-rank shift), is shown in the main text (Figure 2a) and is not repeated here. For each remaining setting, methods are ranked by dataset-weighted excess over the baseline (training-set mean for Wei; mean or additive baseline for Ahlmann-Eltze) on all perturbations versus on the specific subset; methods rising or dropping ≥ 3 ranks are highlighted (green, rises on the specific subset; red, drops). The magnitude of the reshuffle is smaller than in Wei genetic single across all seven settings (maximum shift ≤ 3 ranks), and the top-ranked method changes in four of the seven.

**Figure S6:**
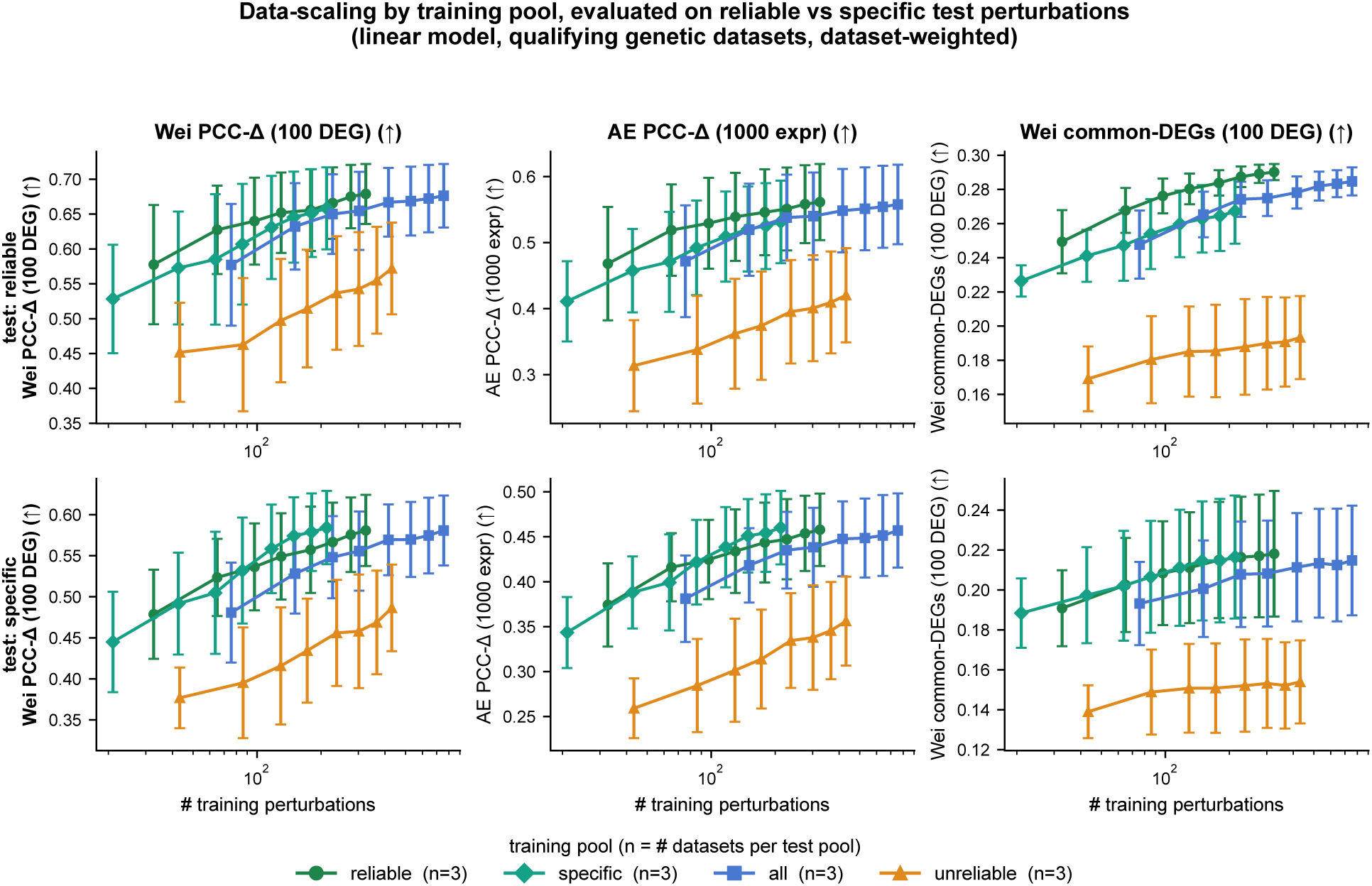
Data-scaling by training pool. Linear-model performance versus the number of training perturbations for four training pools (reliable / specific / all / unreliable), evaluated on reliable (top row) and specific (bottom row) held-out test perturbations, across three evaluation metrics (raw values). Each arm is swept by subset size and dataset-weighted; the legend gives the number of qualifying genetic datasets contributing to each arm. All pools improve with more training data; the reliable, specific and all pools sit above the unreliable pool on both test sets.

**Figure S7:**
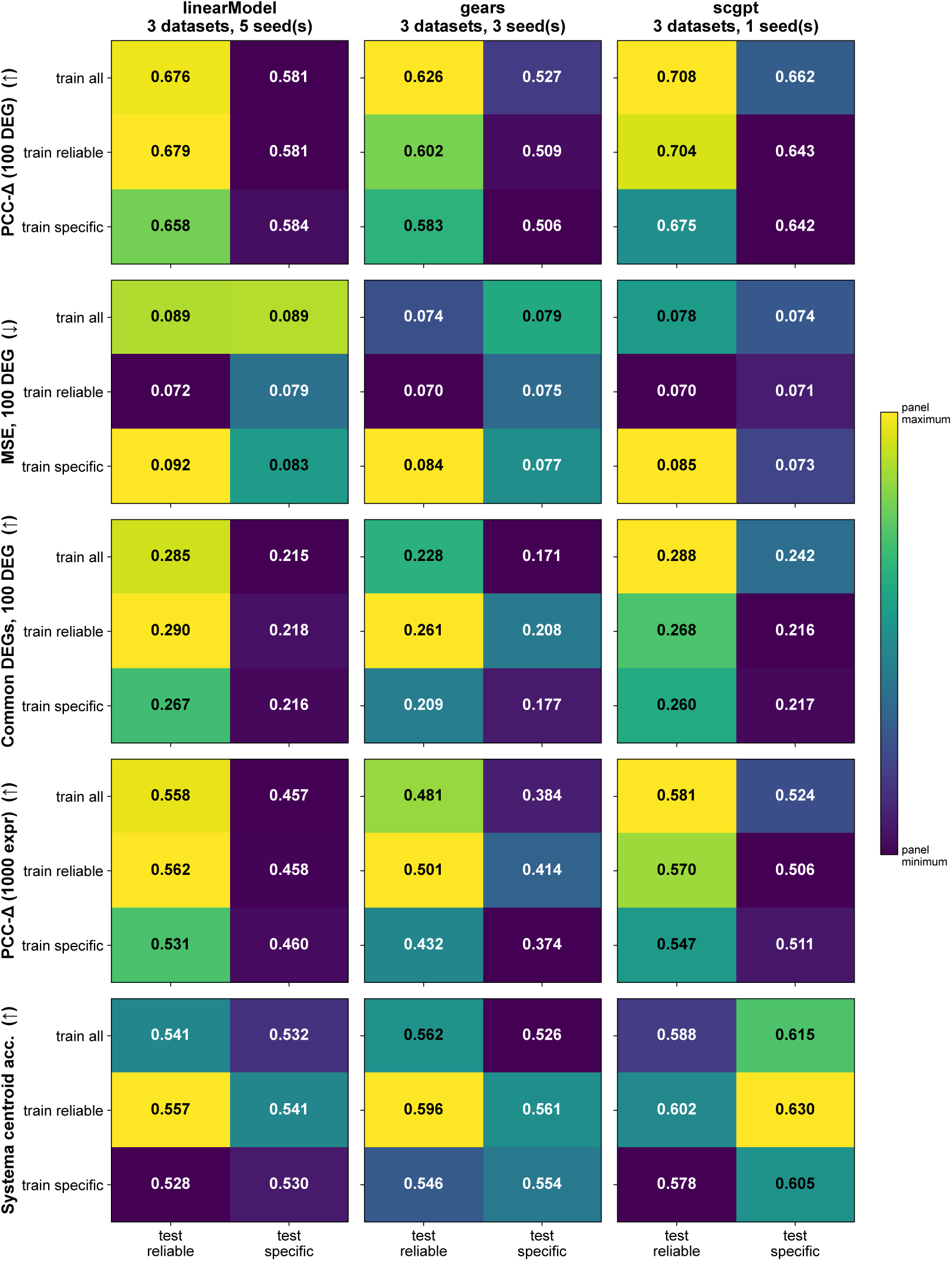
Multi-model retraining grid. Mean performance for each training arm (all / reliable / specific) × test pool (reliable / specific), across five evaluation metrics (rows) and three models (columns: linearModel, GEARS, scGPT), on three genetic datasets. Cell color and annotation give the mean value. Across all three models, training on the reliable subset broadly tracks the performance of training on all perturbations, while training on the specific-only subset is consistently worse on the reliable test pool and no better than the reliable arm on the specific test pool.

**Figure S8:**
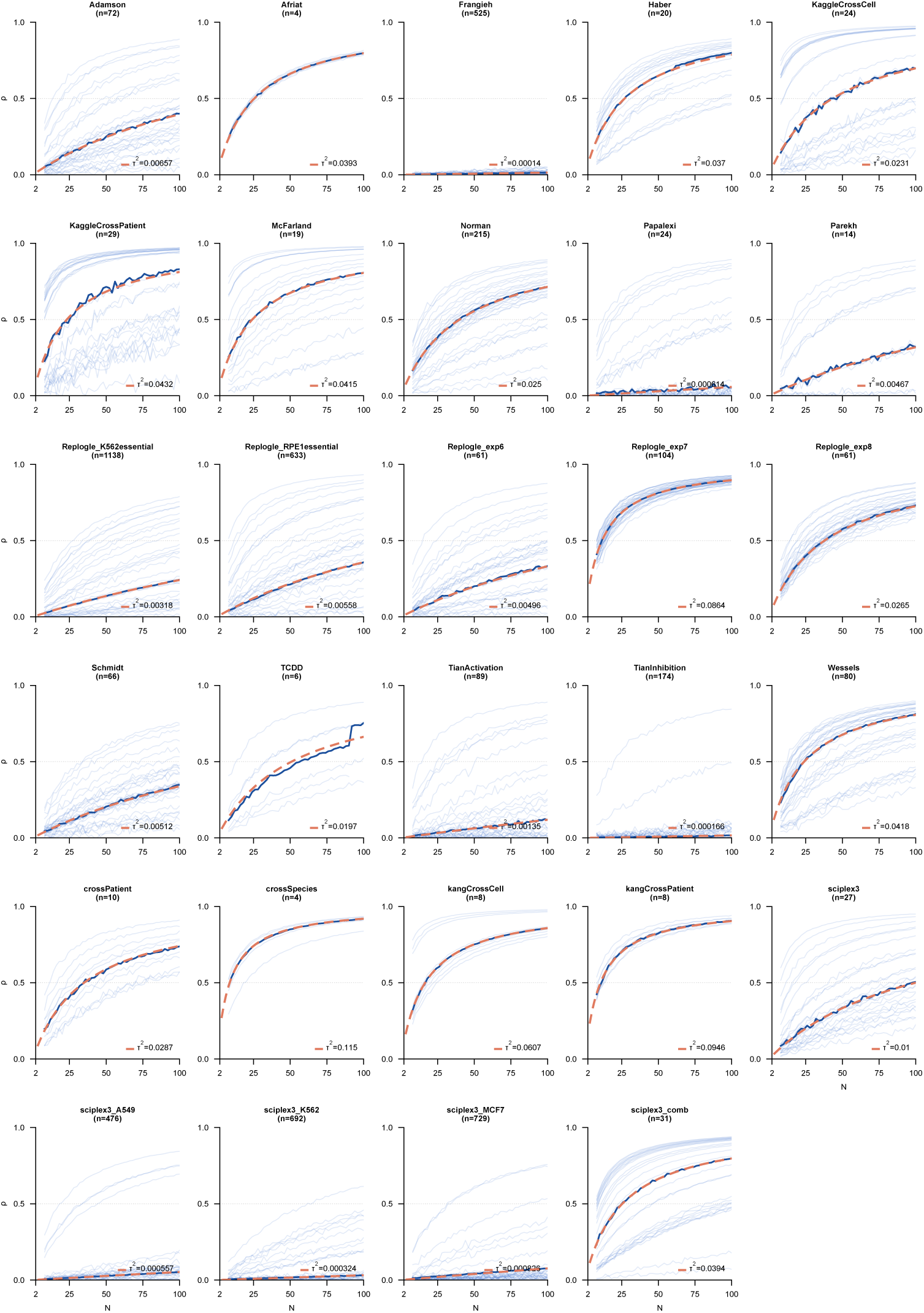
Per-dataset parametric reliability fits. For each dataset, per-perturbation reliability vs. cells per perturbation curves (faint lines), the per-dataset median (solid line), and the parametric *ρ*(*N*) = *Nτ* ^2^*/*(*Nτ* ^2^ + 1) at the median per-perturbation *τ* ^2^ (dashed orange).

